# Multi-functional antibodies are induced by the RTS,S malaria vaccine and associated with protection in a phase I/IIa trial

**DOI:** 10.1101/851725

**Authors:** Liriye Kurtovic, Tanmaya Atre, Gaoqian Feng, Bruce D. Wines, Jo-Anne Chan, Michelle J. Boyle, Damien R. Drew, P. Mark Hogarth, Freya J. I. Fowkes, Elke S. Bergmann-Leitner, James G. Beeson

## Abstract

**Background:** RTS,S is the leading malaria vaccine candidate, but only confers partial efficacy against malaria in children. RTS,S is based on the major *Plasmodium falciparum* sporozoite surface antigen, circumsporozoite protein (CSP). The induction of anti-CSP antibodies is important for protection, however, it is unclear how protective antibodies function.

**Methods:** We quantified the induction of functional anti-CSP antibody responses in healthy malaria-naïve adults (N=45) vaccinated with RTS,S/AS01. This included the ability to mediate effector functions via the fragment crystallizable (Fc) region, such as interacting with human complement proteins and Fcγ-receptors (FcγRs) that are expressed on immune cells, which promote various immunological functions.

**Results:** Our major findings were i) RTS,S-induced antibodies mediate Fc-dependent effector functions, ii) functional antibodies were generally highest after the second vaccine dose; iii) functional antibodies targeted multiple regions of CSP, iv) participants with higher levels of functional antibodies had a reduced probability of developing parasitemia following homologous challenge (p<0.05); v) non-protected subjects had higher levels of anti-CSP IgM.

**Conclusions:** Our data suggests a role for Fc-dependent antibody effector functions in RTS,S-induced immunity. Enhancing the induction of these functional activities may be a strategy to improve the protective efficacy of RTS,S or other malaria vaccines.

## BACKGROUND

After progress in reducing the burden of malaria in the past decade, the annual estimate of *Plasmodium falciparum* malaria cases has plateaued at ∼200 million, which highlights the need for new anti-malarial interventions such as an effective vaccine [1]. RTS,S is the leading malaria vaccine candidate, although efficacy against malaria is only ∼30-50% in young children, and relatively short-lived [2-6]. This is suboptimal compared to licensed vaccines against other pathogens [7, 8], and below the goal set by World Health Organisation and funding partners to develop next-generation vaccines at least 75% efficacious against clinical malaria [9]. Greater efficacy might be achieved by modification of RTS,S or developing new vaccines [10]. Considering RTS,S has been subject to extensive clinical testing and can be safely co-administered with other vaccines as part of standard childhood immunisation programs [3, 11-14], working towards improving RTS,S is an attractive option.

RTS,S targets sporozoites and therefore aims to prevent the initial asymptomatic stage of infection, and the subsequent development of clinical illness, and transmission to mosquitoes [15]. The vaccine is a virus-like particle expressing a fusion protein formed by the hepatitis B surface antigen and the major sporozoite surface antigen, circumsporozoite protein (CSP), administered with AS01 adjuvant [16]. The vaccine includes only the central repeat region (tandem repeat of NANP) and C-terminal region of CSP.

A tangible solution to improve vaccine efficacy is to improve the induction of functional immune responses that mediate protection, since RTS,S/AS01 already achieves very high IgG levels following a 3-dose vaccination schedule [2]. However, the mechanisms of immunity induced by RTS,S and other CSP-based vaccines are unclear, hindering the development of strategies to improve RTS,S or design next-generation vaccines. RTS,S primarily induces antibody and CD4+ T cell responses [17], and a higher magnitude of antibodies to the central repeat region has been associated with protection in some trials [6, 18, 19], and higher avidity IgG to the C-terminal region was associated with higher vaccine efficacy in the phase III trial [20]. However, antibody concentration is only a broad measure that is not strongly indicative of functional activity.

Antibodies can act directly against the target cell, or mediate effector functions via the fragment crystallizable (Fc) region. Antibodies can interact with serum complement protein, C1q, which activates the antibody-dependent classical complement pathway [21]. Additionally, antibodies can engage and cross-link Fcγ-receptors (FcγR) expressed by immune cells to mediate opsonic phagocytosis, or direct killing via antibody-dependent cellular cytotoxicity [22]. These include activating receptors FcγRIIa that is widely expressed on monocytes, macrophages and neutrophils, and FcγRIII that is mostly expressed on neutrophils and natural killer (NK) cells [22]. Although prior studies have shown that antibodies to CSP and the central repeat region can directly inhibit hepatocyte invasion by sporozoites [23-25], there have been few studies of Fc-dependent antibody effector functions against sporozoites.

We previously showed that antibodies can recruit and activate serum complement proteins, which consequently inhibited sporozoite motility and led to cell death [26, 27]. Interestingly, naturally-acquired antibodies to CSP could fix C1q, and children with high levels of these complement-fixing antibodies had a significantly reduced risk of malaria [27]. Furthermore, functional complement-fixing antibodies were induced by RTS,S vaccination in young children in Mozambique [28]. Additionally, we identified specific roles for immune cells, particularly neutrophils, in opsonic phagocytosis of *P. falciparum* sporozoites via interactions with FcγRIIa and FcγRIII (Feng et al., in preparation). Prior studies reported that opsonic phagocytosis activity measured using the promonocytic cell line THP-1 was not a predictor of protection from experimental infection in malaria-naïve volunteers; however, this cell line does not represent all FcγR-mediated functions, and the expression and functions of FcγRs vary across different cell types [22]. Conducting large-scale functional assays with viable *P. falciparum* sporozoites is not feasible, as they cannot be readily cultivated in standard laboratories or obtained in large numbers. To address these technical limitations, we developed high-throughput, cell-free assays that quantify the ability of anti-CSP antibodies to fix human C1q (and other downstream complement components), or interact directly with dimeric recombinant soluble FcγR complexes as a functional surrogate [27, 29, 30]. Furthermore, we used new approaches to measure opsonic phagocytosis activity with fluorescently-labelled beads coated in CSP, and we have previously shown that phagocytosis of CSP-coated beads correlates with phagocytosis of whole sporozoites (Feng et al., in preparation).

Here we quantified anti-CSP antibodies that interact with C1q and FcγRs using samples from a phase I/IIa clinical trial RTS,S/AS01 conducted in healthy malaria-naïve adults. We evaluated the relationship between functional antibodies and protection against controlled human malaria infection (CHMI) challenge.

## METHODS

Detailed materials and methods can be found in the Supplementary.

### Study participants and ethics approval

Healthy malaria-naïve adults were administered three doses of RTS,S/AS01 at month 0, 1 and 2 (ClinicalTrials.gov registry number NCT00075049), and two weeks later underwent homologous CHMI challenge via infectious mosquito bites. Those who remained parasitemia free during the 28 day follow up were considered protected, as previously described [31]. We tested serum samples collected at baseline on day 0 (D0, n=45), and 2-4 weeks post-vaccine dose (PV) 1 (n=45), 2 (n=43) and 3 (n=33). We additionally conducted functional assays using four pools (each pool, n=8) from individuals who were either not-protected (NP1, NP2) or protected (P3, P4) against CHMI challenge.

All study participants had previously provided consent for future use of the samples for research, and ethics approval was also obtained by the Alfred Human Research Committee.

### Antigens

The following antigens used in this study were based on *P. falciparum* 3D7: CSP, peptides representing the central repeat (NANP) [27] and C-terminal (Pf16) [32] regions of CSP, and for some experiments a recombinant construct of the C-terminal region was used (CT) [28].

### Experiments

Total IgG and IgM were measured by standard enzyme-linked immunosorbent assay (ELISA), and functional complement-fixation and FcγR binding assays were conducted as previously described [27, 29, 33]. Opsonic phagocytosis of CSP-coated beads by isolated neutrophils was conducted as previously described (Feng et al., in preparation), and the percentage of cells containing fluorescent positive beads was evaluated by flow cytometry, and presented as phagocytosis index (PI).

### Statistical analysis

Statistical analysis was performed using GraphPad Prism 7.

## RESULTS

### Induction of IgG and IgM in malaria-naïve adults by the RTS,S vaccine

Healthy malaria-naïve adults (N=45) were vaccinated with RTS,S, and serum collected at baseline (day 0, D0) and after each vaccine dose (post-vaccine dose, PV) were tested for antibody reactivity to full length CSP. Anti-CSP IgG significantly increased after the first vaccine dose, and increased further after the second (OD median [IQR]: D0=0.034[0.048], PV1=0.520[0.985], PV2=3.102[0.377]; p<0.001 for both tests). There was some decline in reactivity by the third dose compared to the second (OD median [IQR]: PV3=2.792[0.579]; p=0.029) (**Figure 1A**). To identify which regions of CSP were targeted by antibodies, we tested serum collected at the final time-point for IgG against two peptides representative of the central repeat and C-terminal regions of CSP (NANP and Pf16, respectively). We compared IgG responses from the day of challenge among those who were not-protected (n=17) and protected (n=16) against developing blood-stage parasitemia. IgG reactivity to full-length CSP and Pf16 were not significantly different between groups, but anti-NANP IgG was significantly higher among protected individuals (p=0.363, p=0.136 and p=0.028, respectively) (**Figure 1B**).

**FIGURE 1.**
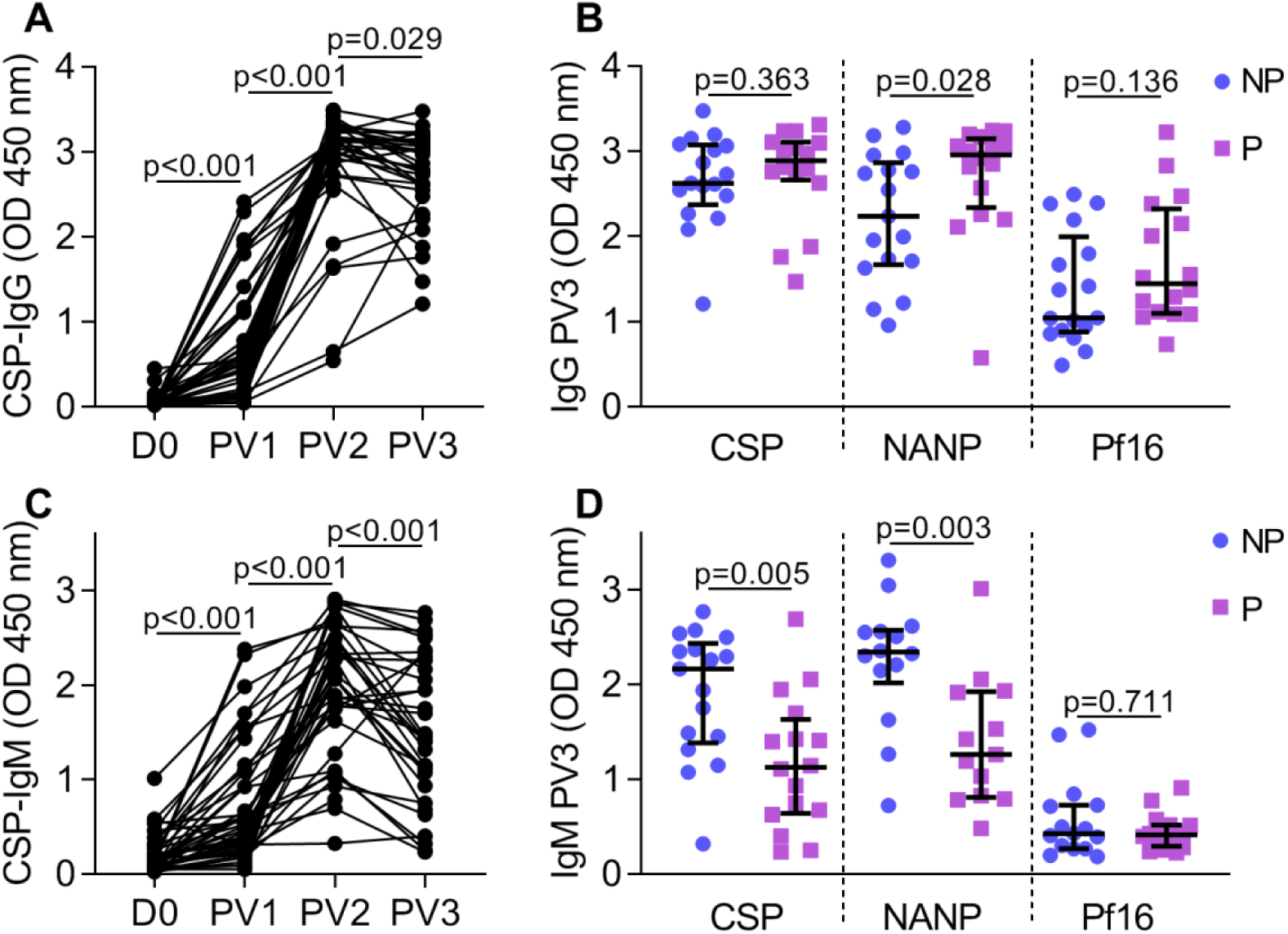
RTS,S-induced IgG and IgM target the central and C-terminal regions of CSP. Serum from adults vaccinated with RTS,S collected at baseline (D0, n=45), and post-vaccine dose 1 (PV1, n=45), 2 (PV2, n=43) and 3 (PV3, n=33) were tested for IgG **(A)** and IgM **(C)** to full length CSP, and those collected PV3 **(B, D)** were also tested against peptides representing the central repeat and C-terminal regions of CSP (NANP and Pf16, respectively). Samples were tested in duplicate and the mean value was graphed, along with the group median and IQR for participants who were not-protected (NP, n=17) and protected (P, n=16) against controlled human malaria infection challenge. Reactivity between paired samples collected at different time points was compared using the Wilcoxon matched pairs signed rank test, and reactivity between unpaired NP and P groups was compared using the and Mann-Whitney U test.

Next, we examined the induction of anti-CSP IgM, which followed a similar pattern to the IgG response in that antibody levels peaked PV2, and declined slightly by PV3 (median [IQR]: D0=0.123[0.187], PV1=0.514[0.710], PV2=2.137[0.876] and PV3=1.454[1.276]) (**Figure 1C**). IgM to CSP and NANP were significantly lower for protected volunteers than not-protected volunteers, whereas anti-Pf16 IgM did not statistically differ between groups (p=0.005, p=0.003 and p=0.711, respectively) (**Figure 1D**).

### Induction of functional antibody responses

We have previously shown that anti-CSP antibodies induced by natural malaria exposure or whole sporozoite immunisation can fix and activate complement, and we also identified antibody-complement interactions as a functional immune mechanism against *P. falciparum* sporozoites [26, 27]. Here we measured the ability of RTS,S-induced antibodies to fix C1q (the first step in classical complement activation) to full length CSP. After two RTS,S vaccinations, strong levels of C1q-fixing antibodies were detected, although responses varied among participants (OD median [IQR]: D0=0[0], PV1=0.010[0.072], PV2=1.095[2.342]; p<0.001) (**Figure 2A**). C1q-fixation responses were significantly lower after the third dose (OD median [IQR]: PV3=0.621[1.327]; p<0.001), but remained substantially higher compared to baseline.

**FIGURE 2.**
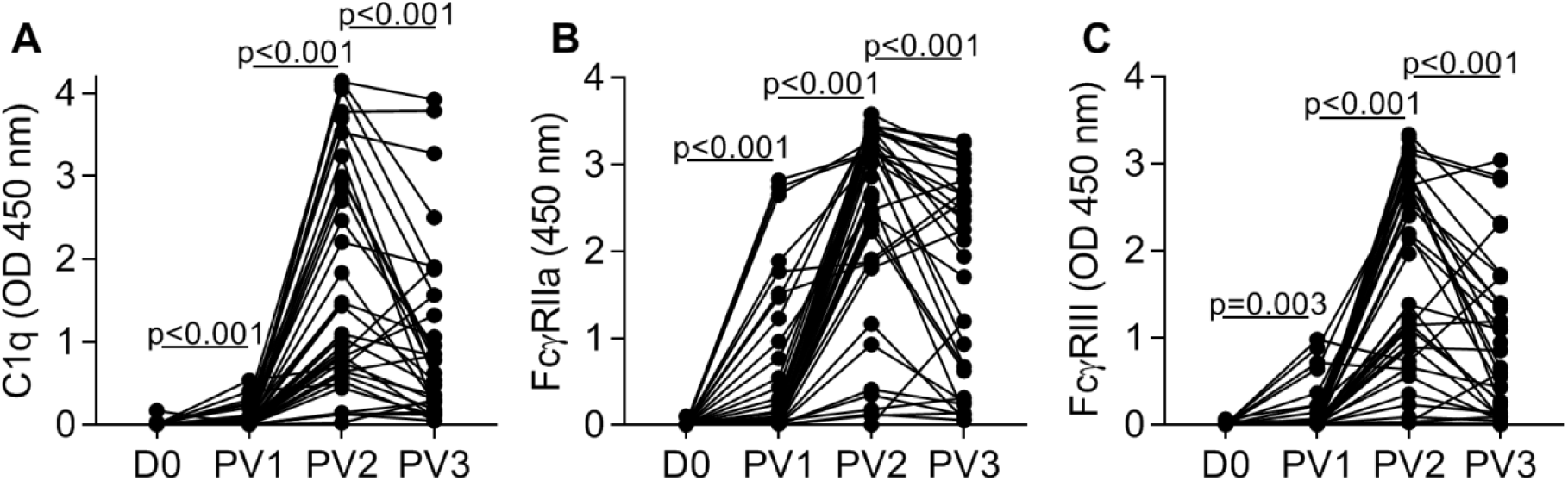
Induction of functional anti-CSP antibodies following RTS,S vaccination. Serum from adults vaccinated with RTS,S collected at baseline (D0, n=45) and post-vaccine dose 1 (PV1, n=45), 2 (PV2, n=43) and 3 (PV3, n=33) were tested for C1q-fixation **(A)**, FcγRIIa **(B)** and FcγRIII **(C)** binding to CSP. Samples were tested in duplicate and the mean value was graphed. Reactivity between paired samples collected at different time points was compared using the Wilcoxon matched pairs signed rank test.

Antibodies can also promote opsonic phagocytosis of *P. falciparum* sporozoites by neutrophils and monocytes, and we recently established that neutrophils play a more prominent role, and involves antibody engagement of FcγRIIa and FcγRIII (Feng et al., in preparation). We measured whether RTS,S-induced antibodies could promote immune complex formation to full length CSP, and interact with dimeric recombinant soluble FcγRs [29]. After the second RTS,S vaccination, there was a strong induction of antibodies that interacted with both FcγRs. Activity was highest after two vaccinations and declined after the third vaccination, but remained above baseline levels (OD median [IQR]: FcγRIIa, D0=0[0], PV1=0.049[0.484], PV2=2.859[1.446], PV3=2.127[2.284]; FcγRIII, D0=0.005[0.014], PV1=0.013[0.058], PV2=1.378[2.007], PV3=0.655[1.431]; p<0.003 for all tests) (**Figures 2B-C**).

### Antibody properties associated with functional activity

We explored the associations between antibody properties and functional activity using data obtained from samples collected after the third RTS,S dose. C1q-fixation and FcγR binding significantly positively correlated with IgG-reactivity to full length CSP, and these associations were slightly stronger when analyses were performed with IgG responses to specific regions of CSP (**Table 1**). To directly confirm the central repeat and C-terminal regions of CSP were targets of functional antibody responses, we tested samples for activity against NANP-repeat peptide and a recombinant construct representing the whole C-terminal region of CSP (CT) instead of the Pf16 peptide; note that IgG to Pf16 and CT strongly correlated (data not shown, Rho=0.850; p<0.001). For these analyses we used samples collected at PV2 due to the limited sample volumes available at the PV3 time-point. Our results demonstrated that antibodies specific to both regions of CSP could interact with C1q and FcγR (**Figure 3**).

**TABLE 1.** Correlations between antibody responses in samples collected PV3 (n=33).

|  |  | IgG |  |  | IgM |  |  | Function (CSP) |  |  |
| --- | --- | --- | --- | --- | --- | --- | --- | --- | --- | --- |
|  |  | CSP | NANP | Pf16 | CSP | NANP | Pf16 | C1q | FcRIIa | FcRIII |
| **IgG** | CSP |  | 0.647 | 0.668 | -0.037 | 0.053 | 0.151 | 0.679 | 0.718 | 0.777 |
|  | NANP |  |  | 0.505 | -0.149 | -0.206 | -0.001 | 0.763 | 0.810 | 0.792 |
|  | Pf16 |  |  |  | 0.096 | 0.165 | 0.410 | 0.719 | 0.752 | 0.813 |
| **IgM** | CSP |  |  |  |  | 0.928 | 0.456 | 0.162 | 0.016 | 0.023 |
|  | NANP |  |  |  |  |  | 0.475 | 0.222 | 0.114 | 0.076 |
|  | Pf16 |  |  |  |  |  |  | 0.204 | 0.254 | 0.192 |
| **Function (CSP)** | C1q |  |  |  |  |  |  |  | 0.893 | 0.913 |
|  | FcRIIa |  |  |  |  |  |  |  |  | 0.940 |
|  | FcRIII |  |  |  |  |  |  |  |  |  |

Spearman's correlation coefficient; significant correlations ( $p < 0.05$ ) were shaded in grey.

**FIGURE 3.**
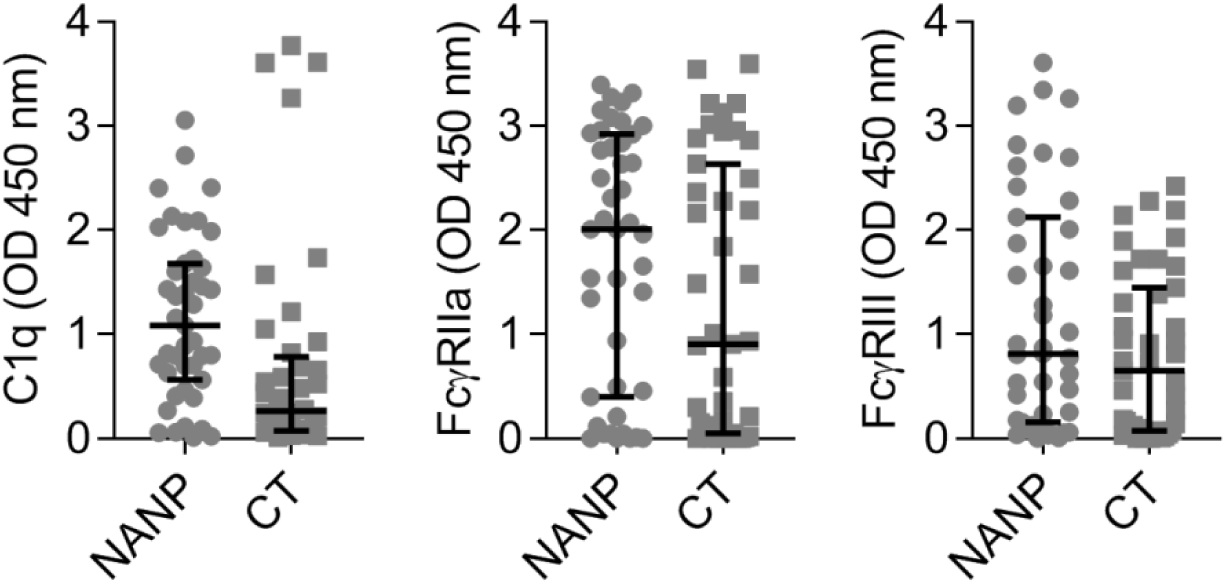
Functional antibody responses to the central repeat and C-terminal regions of CSP. Serum from adults vaccinated with RTS,S collected post-vaccine dose 2 (n=43) were tested for C1q-fixation, FcγRIIa and FcγRIII binding to antigens representing the central repeat and C-terminal regions of CSP (NANP and CT, respectively). Samples were tested in duplicate and the mean value was graphed, along with the group median and IQR.

In contrast, there were no significant correlations between antibody function and IgM (**Table 1**). IgM does not engage FcγR; however, IgM has the potential to strongly activate complement [34], so the lack of association with C1q-fixation was unexpected. It is also noteworthy that IgM did not significantly correlate with IgG. To explore further, we quantified the relative ratio of IgG-to-IgM, and found that while IgM responses were prominent after the first vaccine dose, antibody responses increasingly became IgG-skewed after the second and third vaccinations (**Figure 4A**). All participants had a similar ratio of IgG-to-IgM after the first vaccine dose, but after the second and third dose those who were protected against CHMI had a higher ratio of IgG-to-IgM than not-protected volunteers (**Figure 4B**).

**FIGURE 4.**
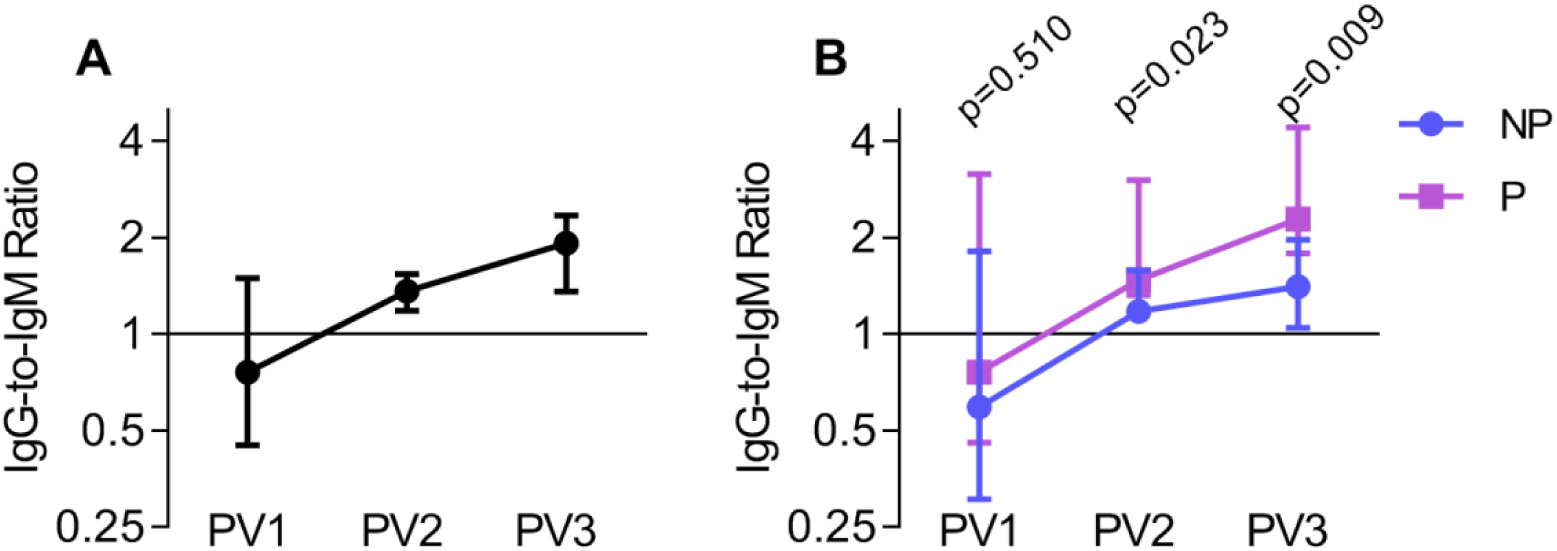
Ratio of IgG-to-IgM after each RTS,S dose. Serum from adults vaccinated with RTS,S collected post-vaccine dose 1 (PV1, n=45), 2 (PV2, n=43) and 3 (PV3, n=33) were tested for IgG and IgM to CSP. The ratio of IgG-to-IgM was calculated at each time-point, and the group median and 95% CI of the median were graphed. **(A)** All participants, and **(B)** those who were not-protected (NP, n=17) and protected (P, n=16) against controlled human malaria infection were shown. The ratio of IgG-to-IgM (for NP and P groups) at all time-points was compared using Mann-Whitney U test.

### Associations between functional antibodies and protection against CHMI

Participants were defined as having high (function score 1) or low (function score 0) activity for each functional antibody type based on whether they were above or below the median (median OD values for each response type were: C1q=0.621, FcγRIIa=2.127 and FcγRIII=0.655) (**Figure 5A**). Participants with high activity for C1q-fixation, or binding to FcγRIIa or FcγRIII were more likely to remain malaria free during the 28 day follow-up than those with low activity (**Figure 5B**). Furthermore, when we examined all three responses together, participants with high activity for one or more functional antibody types had a significantly delayed onset of parasitemia compared to those with overall low activities (total function score of ≥1 versus 0; p=0.026) (**Figure 5C**).

**FIGURE 5.**
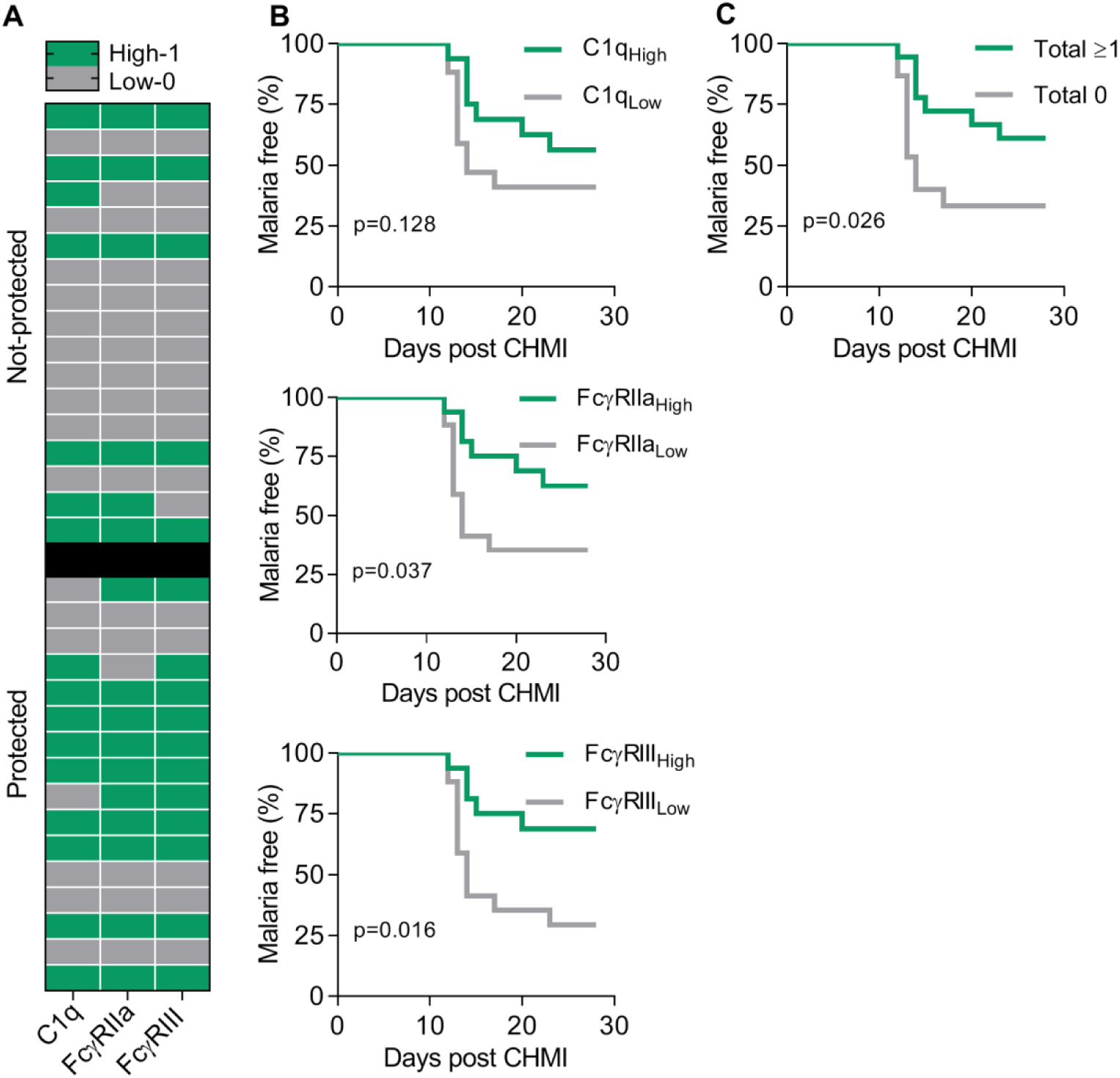
Relationship between functional antibodies and protection against controlled human malaria infection (CHMI). Serum from adults vaccinated with RTS,S collected post-vaccine dose 3 (PV3, n=33) were tested for C1q-fixation, FcγRIIa and FcγRIII binding to CSP. Participants were defined as having high or low activity (function scores of 1 and 0, respectively) for each functional antibody response type based on the median OD values: C1q=0.621, FcγRIIa=2.127 and FcγRIII=0.655. **(A)** Heat-map of participants with high and low antibody activity who were protected or not-protected against CHMI challenge (rows represent the individual subject). **(B)** Kaplan-Meier survival curves of individuals with high and low activity for each functional antibody response type and malaria status during 28 days follow-up, and **(C)** of individuals with high activity for at least one functional antibody response type or low for all functional responses (total function scores of 0 and ≥1, respectively). Kaplan-Meier survival curves were analysed using the Gehan-Breslow-Wilcoxon test.

To confirm that C1q-fixation and FcγR binding assays were indicative of complement activation and opsonic phagocytosis, respectively, we further examined the functional activities of pooled serum samples from individuals vaccinated with RTS,S (each pool, n=8). Two pools contained individuals who were not-protected against CHMI (NP1, NP2), and two pools contained individuals who were protected against CHMI (P3, P4). Notably IgG reactivity to full length CSP was similar among the four pools, whereas IgM reactivity was variable (**Figure 6A**).

**FIGURE 6.**
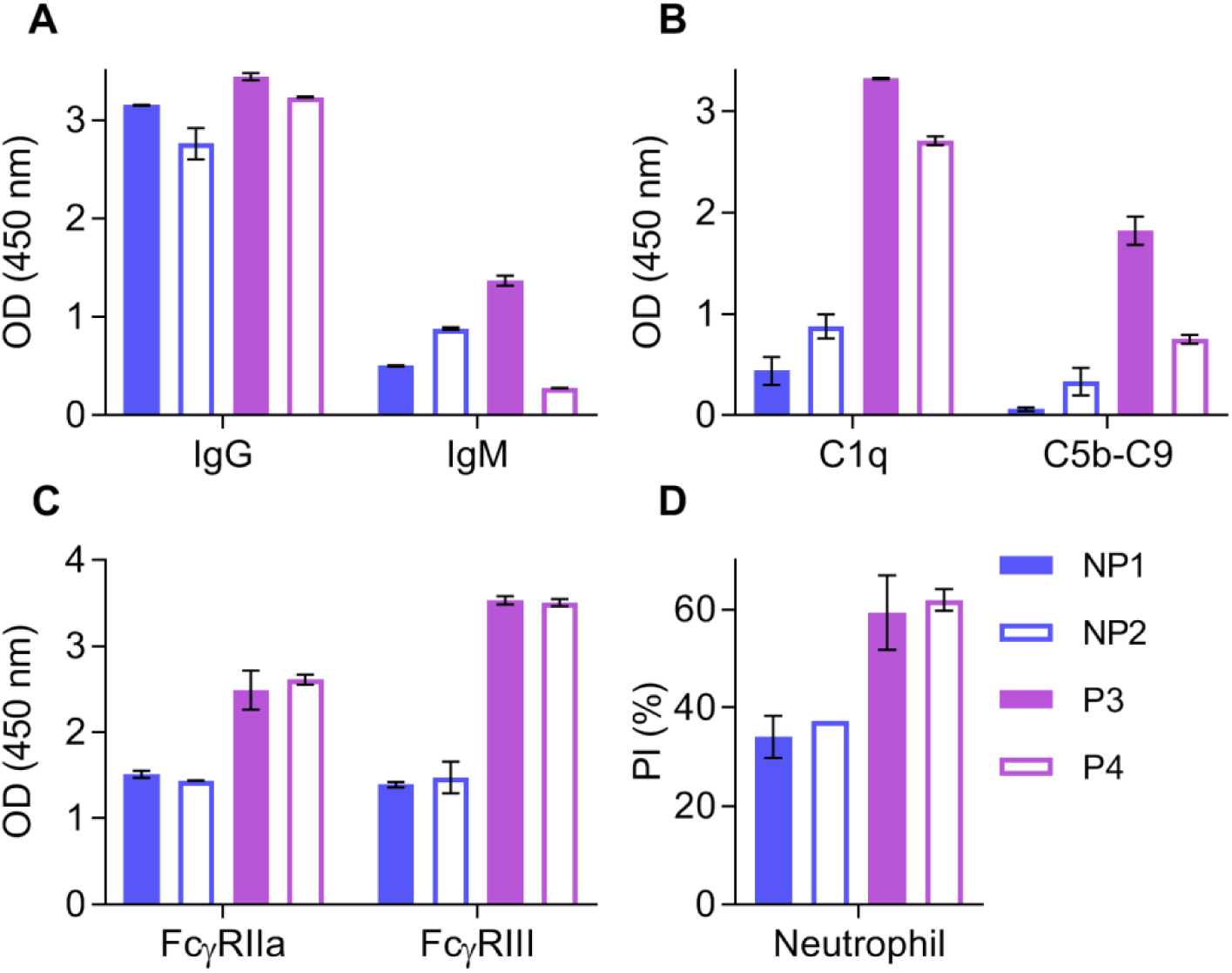
Pooled serum activates complement and promotes opsonic phagocytosis. Pooled samples from individuals vaccinated with RTS,S (each pool, n=8) who were not protected (NP1, NP2) or protected (P3, P4) against CHMI challenge were tested for IgG and IgM reactivity to full length CSP **(A)**, C1q and C5b-C9-fixation to CSP **(B)**, FcγRIIa and FcγRIII binding to CSP **(C)**, and opsonic phagocytosis of CSP-coated beads by freshly isolated neutrophils **(D)**, and. Pools were tested in duplicate, and the mean and range of duplicates were graphed.

We tested each pool for the ability to fix C1q, and determined whether this led to recruitment and activation of downstream complement proteins by measuring the fixation of C5b-C9, which is involved in the terminal phase of complement and forms the membrane attack complex. Indeed, pooled samples could fix C1q and C5b-C9, and responses were substantially higher among protected pools compared to the not-protected pools (C5b-C9-fixation, OD mean [range]: P3=1.822[1.683-1.962], P4=0.749[0.703-0.795], NP1=0.059[0.041-0.077] and NP2=0.331[0.197-0.465]) (**Figure 6B**).

The pools were also used to confirm that RTS,S-induced antibodies could bind FcγRs and promote opsonic phagocytosis (**Figures 6C-6D**). We focussed on neutrophil phagocytosis as our recent studies established that neutrophils are the major cell type in blood that is active in phagocytosis of sporozoites (Feng et al., in preparation). The pools were used to opsonise CSP-coated beads for phagocytic uptake by freshly isolated human neutrophils, and functional activity was greater for the protected pools than the not-protected pools (PI mean [range]: P3=59.35[51.8-66.9], P4=61.9[59.7-66.9], NP1=34.05[29.8-38.3] and NP2=37.3[37.3-37.3], respectively). Additionally, binding to FcγRIIa and FcγRIII was greater for protected pools compared to not-protected pools, which was similarly observed for C1q-fixation.

## DISCUSSION

Knowledge of the immune mechanisms induced by RTS,S, and responses that confer protection is limited, which hinders our capacity to enhance RTS,S or develop new malaria vaccines. Here we evaluated Fc-mediated functional antibody responses induced by RTS,S vaccination in healthy malaria-naïve adults in a phase I/IIa clinical trial. Our major findings were i) RTS,S-induced antibodies could fix and activate complement, bind to FcγRIIa and FcγRIII, and mediate opsonic phagocytosis by neutrophils; ii) functional antibodies were higher after the second vaccine dose rather than the third dose; iii) functional antibodies targeted both central repeat and C-terminal regions of CSP and correlated with IgG reactivity but not IgM, iv) participants with higher levels of functional antibodies had a reduced probability of developing parasitemia following homologous CHMI challenge (p<0.05); v) non-protected subjects had higher levels of IgM reactivity to CSP and a higher IgM-to-IgG ratio. These data give important insights into the mechanisms of RTS,S-induced immunity to inform future malaria vaccine development, and bring us closer to identifying functional immune correlates of protection.

Antibodies to CSP induced by RTS,S were able to fix human complement proteins, including the initiation protein C1q, and downstream proteins C5b-C9. We previously established antibody-complement interactions as an immune mechanism against *P. falciparum* sporozoites [26, 27], and that complement-fixing antibodies could be induced by RTS,S vaccination in young children in Mozambique [28]. However, in that trial, RTS,S was formulated with the previously used AS02 adjuvant. Therefore, our current study specifically demonstrates that RTS,S/AS01 can also induce this functional antibody in malaria-naïve adults. We additionally show that RTS,S-induced antibodies can mediate opsonic phagocytosis by neutrophils. While phagocytosis has been previously reported using the THP-1 promonocyte cell line [35, 36], an advantage of our study was that we measured phagocytic uptake by freshly isolated neutrophils, which are the most abundant leukocyte in the blood, and account for the majority of phagocytosis activity in assays using whole blood (Feng et al., in preparation). Furthermore, vaccine-induced antibodies could directly bind to FcγRIIa and FcγRIII, which are expressed on neutrophils, NK cells and other cells. This has not been previously examined in RTS,S studies, but these interactions play important roles in immunity to other pathogens and appear to be important in immunity in the partially successful RV144 vaccine trial for HIV [30, 37].

Strong levels of functional antibodies were acquired after two RTS,S vaccinations, and there was no increase after the third vaccination, instead reactivity had declined. Similarly, a reduced response between the second and third dose was also observed for total IgG and IgM. Considering that peak antibody responses were induced after the second vaccine dose, there may have been no additional benefit to receiving the third vaccination. Indeed other trials in malaria-naïve adults have found similar immunogenicity and efficacy (∼50%) when immunised with two or three RTS,S doses [38, 39]. This contrasts studies of malaria-exposed children, whereby a 3-dose immunisation schedule resulted in enhanced IgG responses compared to a 2-dose immunisation schedule [40]. Furthermore, recent evidence suggests that a delayed fractional (reduced antigen amount) third dose improves efficacy compared to the standard 3 full dose regimen [41], but this is yet to be confirmed in field evaluations [42]. Clearly there are important differences in immunogenicity and efficacy following vaccination with 2 or 3-dose regimens of RTS,S, particularly among children and adults, but in this study we report high immunogenicity after two vaccine doses.

A key finding was that participants who acquired high levels of anti-CSP antibodies that fixed C1q or bound FcγRs generally had a reduced probability of malaria following CHMI. Importantly, participants with high activity for at least one functional antibody response type had a significantly delayed onset of parasitemia than those with overall low functional activities. Among serum pools from vaccinated individuals, complement and opsonic phagocytosis functional activities were substantially higher for the protected pools compared to the not-protected pools. Since this was a phase I/IIa challenge trial the number of study subjects was relatively small, which limits the statistical power to further investigate associations with protection. Further studies in larger RTS,S challenge trials and field-based trials in target populations are needed to confirm protective associations identified in this study, and to define correlates of protection.

The relationship we observed between higher functional antibody responses and protection is in some agreement with other published studies. Indeed, high levels of naturally-acquired anti-CSP antibodies that fix complement have been associated with a reduced risk of malaria [27]. Opsonic phagocytosis by THP1 cells have not been consistently associated with protection in phase I/IIa CHMI trials [35, 36]. Our recent studies demonstrated that sporozoites are mostly phagocytosed by neutrophils in whole blood assays, and there are important differences in FcγR expression and function between THP-1 cells and neutrophils (Feng et al., in preparation). In the current study, when using freshly isolated human neutrophils we found a clear differentiation between protected and not-protected pools. Another novel aspect was that we measured binding to FcγRIII, which is typically poorly expressed on THP-1 cells and resting monocytes [43], but is expressed on resting neutrophils and NK cells [22], and FcγRIII binding strongly correlated with NK cell activation against non-malaria pathogens in vitro [44, 45]. Interestingly, there was a more pronounced distinction between time to becoming malaria positive and high activity for binding to FcγRIII than FcγRIIa. This receptor may also be important as Kupffer cells, the resident liver macrophage, have also been shown to express FcγRIII in addition to FcγRIIa [46, 47], and may clear opsonised sporozoites [48]. It is noteworthy that FcγRIII exists as two forms; FcγRIIIa has a transmembrane domain whereas FcγRIIIb has a glycosylphosphatidylinositol anchor. FcγRIIIa is expressed on NK cells, neutrophils and macrophages, whereas FcγRIIIb is expressed on neutrophils. The dimeric FcγRIII binding assay used was a measure of pan FcγRIII binding activity [22].

Fc-dependent responses to full length CSP correlated strongly with IgG to the central repeat and C-terminal regions of the protein, and we directly showed that antibodies to both regions of CSP could bind FcγRIIa and FcγRIII. The relative contribution of antibodies to each epitope in relation to functional activity should be further investigated, particularly because antibody levels to the central repeat region are generally higher in protected volunteers in other RTS,S vaccine trials [6, 18, 19]. Generally, field evaluations only measure antibodies to the R32LR peptide representing the central repeat region, and overlook the potential role of C-terminal antibodies [49]. However, a recent study was the first to report that RTS,S-induced antibodies to both regions of CSP of a specific IgG subclass were associated with protection against clinical malaria in children [50]. We also measured the induction of anti-CSP IgM, which had no correlation with total IgG responses. Interestingly, protected volunteers had significantly less IgM than not-protected volunteers, which was similarly observed in another RTS,S vaccine study of malaria-exposed children [50], and tended to have higher IgG-to-IgM ratios compared to not-protected volunteers. While IgM does not interact with FcγR it can potently activate complement; however, IgM responses had no significant correlation with any functional antibody responses. It may be that the higher IgM levels inhibit FcγR functions involved in protection, or IgM might be reflective of other differences in immune function between groups. This observation warrants investigation in future studies.

In summary, we demonstrated the induction of antibodies that interact with complement and FcγRs following RTS,S vaccination in healthy malaria-naïve adults. Participants who acquired greater levels of these antibodies generally had a reduced probability of infection or delayed onset of parasitemia following CHMI challenge. These data provide promising evidence that Fc-dependent effector functions may play a role in vaccine-induced immunity, and encourage further investigation in larger RTS,S trials, particularly in malaria-exposed cohorts. This study brings us closer to identifying immune correlates of protection, which will be crucial for developing highly efficacious vaccines. Such knowledge may shed light on why RTS,S is only partially efficacious, provide insight into how RTS,S could be modified to enhance efficacy in target populations, and aid the development of highly efficacious malaria vaccines.

## NOTES

## Acknowledgements

We thank all study participants and staff at Walter Reed Army Institute of Research (WRAIR) who were involved in the clinical trial. We also thank Elizabeth Duncan for administrative support. The Mal-027 clinical trial (NCT00075049) was sponsored by the US Army Medical Research and Development Command in collaboration with WRAIR and GlaxoSmithKline Biologicals SA (GSK). RTS,S is a malaria vaccine candidate developed by GSK and GSK was provided the opportunity to review a preliminary version of this manuscript for factual accuracy, but the authors are solely responsible for final content and interpretation

## Conflict of interest

The authors declare they have no competing interests.

## Funding

This work was funded by the National Health and Medical Research Council (NHMRC) of Australia (Program Grant and Project Grant to JGB; Project Grant to PMH and BDW; Senior Research Fellowship to JGB; Career Development Fellowship to FJIF), National Institutes of Health, the Military Infectious Disease Research Program (MIDRP, TA and EBL), Australian Government Research Training Program Scholarship to LK, Monash Postgraduate Publication Award to LK, and the Australian Society for Parasitology Network Researcher Exchange, Training and Travel award to LK. The Burnet Institute is supported by a Victorian State Government Operational Infrastructure Support grant, and the NHMRC Independent Research Institutes Infrastructure Support Scheme. JGB, FJIF, LK, MJB, JAC, and GF are supported by the Centre for Research Excellence in Malaria Elimination, which is funded by the NHMRC, Australia.

## Presentation of the data

A selection of data from this manuscript has been presented at the following meetings: Malaria Gordon Research Conference (July 2019), Les Diablerets, Switzerland; Australian Society for Parasitology (July 2019), Adelaide, Australia; Malaria in Melbourne (October 2019), Melbourne, Australia.

## SUPPLEMENTARY METHODS

### Study participants and ethics approval

Serum samples from individuals immunised with RTS,S/AS01 were obtained from a previously conducted randomised control phase I/IIa clinical trial (ClinicalTrials.gov registry number NCT00075049). Healthy malaria-naïve adults were administered three doses of RTS,S/AS01 at month 0, 1 and 2 of the study, and two weeks later underwent homologous CHMI challenge via infectious mosquito bites. Those who remained parasitemia free during the 28 day follow up were considered protected, as previously described [31]. We tested serum samples collected at baseline on day 0 (D0, n=45), and 2-4 weeks post-vaccine dose (PV) 1 (n=45), 2 (n=43) and 3 (n=33). We additionally conducted functional assays using four pools (each pool, n=8). Pooled samples contained serum from individuals collected after RTS,S vaccination who were either protected or not-protected against CHMI challenge, and were defined as not-protected 1, not-protected 2, protected 3 and protected 4 (NP1, NP2, P3, P4, respectively). Serum samples were heat-inactivated at 56°C for 45 minutes to inactivate heat-sensitive complement proteins prior to use.

All study participants had previously provided consent for future use of the samples for research. Consent to publish was not required as the samples were de-identified. The study was deemed exempt from human use (WRAIR#2142) and ethics approval was also obtained by the Alfred Human Research Committee (The Alfred, Melbourne). The investigators have adhered to the policies for protection of human subjects as prescribed in AR 70-25.

### Antigens

The following antigens used in this study were based on *P. falciparum* 3D7: recombinant CSP expressed in *Pichia pastoris* (Sanaria, Rockville, USA), synthetic (NANP)_15_ peptide representing the central repeat region of CSP (NANP, Life Tein, Hillsborough, USA) [27], and synthetic peptide representing a portion of the C-terminal region of CSP (Pf16, sequence: EPSDKHIKEYLNKIQNSLSTEWSPCSVTCGNGIQVRIKPGSANKPKDELDYANDIEKKI CKMEKCS) [32]. In functional antibody experiments, we alternatively used a recombinant construct representing the entire C-terminal region of CSP expressed in the mammalian cell line HEK293 (CT) [28].

### Antibody detection by enzyme-linked immunosorbent assay (ELISA)

Total IgG and IgM were measured by standard ELISA as follows: 96-well flat bottom MaxiSorp plates (Thermo Fisher Scientific, Waltham, USA) were coated in 0.5 μg/ml antigen in PBS at 4°C, and then blocked using 1% casein in PBS at 37°C. Serum samples were tested in duplicate at 1/8000 and 1/500 dilutions (for IgG and IgM, respectively) in 0.1% casein in PBS. Bound antibodies were detected using goat anti-human IgG and IgM conjugated to horse radish peroxidase (HRP, Millipore, Burlington, USA), both tested at 1/2500 dilution in 0.1% casein in PBS. Finally, plates were incubated with tetramethylbenzidine (TMB) substrate (Thermo Fisher Scientific) for 10 and 15 minutes to detect IgG and IgM, respectively. The reaction was stopped using 1M sulfuric acid and optical density (OD) was immediately measured at 450 nm. Note that for all plate-based assays, between each step plates were washed thrice in PBS Tween20 0.05%, and all incubations were conducted at room temperature unless specified otherwise.

### Complement-fixation assay

The complement-fixation assay was conducted as previously described [27, 33]. Briefly, plates were coated and blocked as described in the standard ELISA protocol, and serum samples were tested in duplicate at 1/100 dilution. Plates were then incubated with purified human C1q (10 μg/ml, Millipore), and C1q-fixation was detected using rabbit anti-C1q IgG (in-house) followed by goat anti-rabbit IgG conjugated to HRP (Millipore), both tested at 1/2000 dilution. Reagents were prepared in 0.1% casein in PBS, and reactivity was measured after 12 minutes of TMB substrate incubation. To measure the terminal phase of complement activation, we adapted the C1q-fixation assay to measure downstream complement proteins, C5b-C9, which form the membrane attack complex, as previously described [27].

### FcγR binding assay

The FcγR binding assay was conducted as previously described [29]. Briefly, plates were coated as described in the standard ELISA protocol. Plates were blocked with 1% bovine serum albumin (BSA) in PBS, and serum samples were tested in duplicate at 1/250 dilution. Plates were then incubated with biotin-labelled dimeric recombinant soluble FcγRIIa (0.2 μg/ml) and FcγRIII (0.1 μg/ml), and FcγR binding was detected using high sensitivity streptavidin conjugated HRP (Thermo Fisher Scientific) at 1/10000 dilution; both steps were performed at 37°C. Reagents were prepared in 1% BSA in PBS, and reactivity was measured after 7 and 15 minutes of TMB substrate incubation for FcγRIIa and FcγRIII, respectively.

### Functional opsonic phagocytosis assay

Opsonic phagocytosis of CSP-coated beads by isolated neutrophils was conducted as previously described (Feng et al., submitted). Neutrophils were isolated from peripheral blood from healthy Melbourne donors using Ficoll gradient centrifugation, followed by dextran segmentation and hypotonic lysis. The isolated neutrophils were re-suspended in RPMI-1640 supplemented with 10% fetal bovine serum and 2.5% heat-inactivated human serum, and kept on ice until use. CSP-coated beads (5×10^7^ beads/ml) were opsonised for 1 hour with serum pools at 1/160, washed thrice with RPMI-1640, and then incubated with neutrophils for 20 minutes at 37°C to allow phagocytosis to occur. Samples were then washed with PBS-new born calf serum 1% via centrifugation at 300 g for 4 minutes, and the percentage of cells containing fluorescent positive beads was evaluated by flow cytometry, and presented as phagocytosis index (PI).

### Standardisation

For plate-based assays all samples were tested in duplicate, and those with high variability (>25%) were re-tested or excluded (unless OD difference <0.1) to ensure accurate results. Raw data were subtracted from no-serum negative control wells, and standardised for plate-to-plate variability using positive controls that were included on each plate.

### Statistical analysis

The following non-parametric tests were conducted where appropriate (GraphPad Prism 7): Wilcoxon matched pairs signed rank test for paired samples, Mann-Whitney U test for unpaired samples, and Spearman’s correlation coefficient (Rho). To evaluate the relationship between functional antibody responses and time to becoming malaria positive, we plotted Kaplan-Meier survival curves of participants defined as having high or low activity based on the median OD for each functional response (C1q, FcγRIIa and FcγRIII). Groups were compared using the Gehan-Breslow-Wilcoxon test.

